# BeamDelta: simple alignment tool for optical systems

**DOI:** 10.1101/833368

**Authors:** Nicholas J. Hall, David Miguel Susano Pinto, Ian M. Dobbie

## Abstract

BeamDelta is a tool to help align optical systems. It greatly assists in assembling bespoke optical systems by providing a live view of the current laser beam position and a reference position.

Even a simple optical setup has multiple degrees of freedom that affect the alignment of beam paths. These degrees of freedom rise exponentially with the complexity of the system. The process of aligning all the optical components for a specific system is often esoteric and poorly documented, if it is documented at all.

Alignment methods used often rely on visual inspection of beams impinging on pinholes in the beam path. Typically requiring an experienced operator staring at diffuse reflections for extended periods of time. This can lead to a decline in accuracy due to eye strain, flash blindness as well as symptoms such as headaches and, possibly, more serious retinal damage.

Here we present BeamDelta a simple alignment tool and accompanying software interface which allows users to obtain accurate alignment as well as removing the necessity of staring at diffuse laser reflections. BeamDelta is a robust alignment tool as it doesn’t require any precise alignment itself.

## Introduction

Many laboratories rely on microscopes for their ongoing research. However, commercial offerings often lag behind recent developments in the field or they might not be tailored to specific user’s needs. As such, there often arises a need for laboratories to build their own bespoke microscopy systems. Many bespoke optical systems are highly complex and delicate systems requiring many, sometimes several dozen, optical components. This is especially true of cutting edge microscopy setups such as the Lattice Light Sheet[1], 4*π* whole cell single molecule switching[2], iSIM[10], or Adaptive Optical SIM system[11, 9, 8]

Often, even within the group which build such instruments, the alignment protocol is ill defined and imprecise. Typically, systems are initially aligned with a small beam and no lenses. Pinholes or irises are placed at critical points in the optical system and then a light beam, usually a laser with a low divergence angle, is aligned to these points. Lenses are then added one at a time. Once a lens is added, propagation of the beam is checked further along the optical path to ensure the beam is passing through the centre of the lens and the lens angle is judged by ensuring the back reflections are centred on the incident beam. This process is repeated for each lens. The quality of the alignment depends upon significant expertise and subjective judgement to both see how well aligned the beam is and to decide when it is sufficiently well aligned to move on to the next element in the system.

This approach has two obvious problems. The first is that the human eye has limited resolving power of approximately 3 to 6 arcseconds [5]. At a distance of 50 cm, this is a resolution limit of approximately 10 µm. This doesn’t account for parallax errors which can further reduce the resolution and introduce systematic errors, nor does is account for the user’s ability to accurately judge the comparison between where a laser spot currently is and where it was at some point in the past [3]. The second problem is that this method of alignment requires users to stare at diffuse laser reflections for extended periods of time which carries an inherent safety risk [7]. Again, this problem can be exacerbated if a user tries to minimise the parallax errors and get level with the optical axis as they then run the risk of accidentally observing collimated laser light or specular reflections.

Our tool, BeamDelta, aims to deal with both of these problems. It measures the position of the laser beam and provides real time feedback on its distance from an aligned position. The position is measured from a spot centroid in a camera image. The measured position can be significantly better than the optical resolution or pixel size, 5–10 µm with modern low cost cameras [4], and easily beat what is achievable by the human eye, without the parallax error. Additionally, the user does not need to observe the laser or reflections directly thereby reducing the safety risk of direct exposure of the eye to laser light.

## Methods

### Implementation

There are two aspects to BeamDelta: the hardware and the software.

#### Hardware

In order to describe our implementation and usage of the system in practice, we discus the details of two use cases of BeamDelta for optical system alignment. These cover the addition of a single lens into an existing system and the co-alignment of two beams, e.g. two different wavelength lasers.

#### Use Case 1: Lens Alignment

In this case we are adding a lens to a system which has a small aligned beam already propagating along the expected beam path. The objective is to add an additional lens to the system and achieve good alignment with the lens perpendicular to the beam and centred on it. To perform re-alignment upon addition of a single lens, only a single image is required. So long as the image plane is not at the focus of the lens, then offsets in the beam path from ideal will lead to offsets in the beam position. In this case the easiest approach is to have a compact CMOS camera directly in the beam path. The camera can be attached to a post via a c-mount adaptor and placed directly in the beam path.

#### Use Case 2: Laser Co-alignment

In this use case we already have one laser propagating along the system and we wish to align a second laser so it propagates at the same position and angle as the existing beam. To determine both the position and angle of a beam, two image planes must be observed. We do this by having two cameras mounted in a cage system with a non-polarising 50:50 beamsplitter cube between them. In order to achieve reasonable angular sensitivity the two image planes must be at different optical positions, so different distances from the 50:50 beamsplitter.

Additionally we find it helpful to mount a 45° plane mirror before the beamsplitter cube to allow the whole alignment apparatus to be out of plane of the optical beampath and easily moved to different locations in the system. The apparatus as a whole is then mounted on a post and can be placed in the beampath.

The Dual Camera configuration we use is shown in Figure 1. The lower and upper cameras are at 50 mm and 230 mm from the centre of the beamsplitter respectively, leading to a path difference of 180 mm. The centre of the beamsplitter is 65 mm from the centre of a 45° plane mirror. Having adjustment thumb screws on this 45° mirror enables small adjustments to be easily made to ensure the beams fall on the sensors.

**Figure 1:**
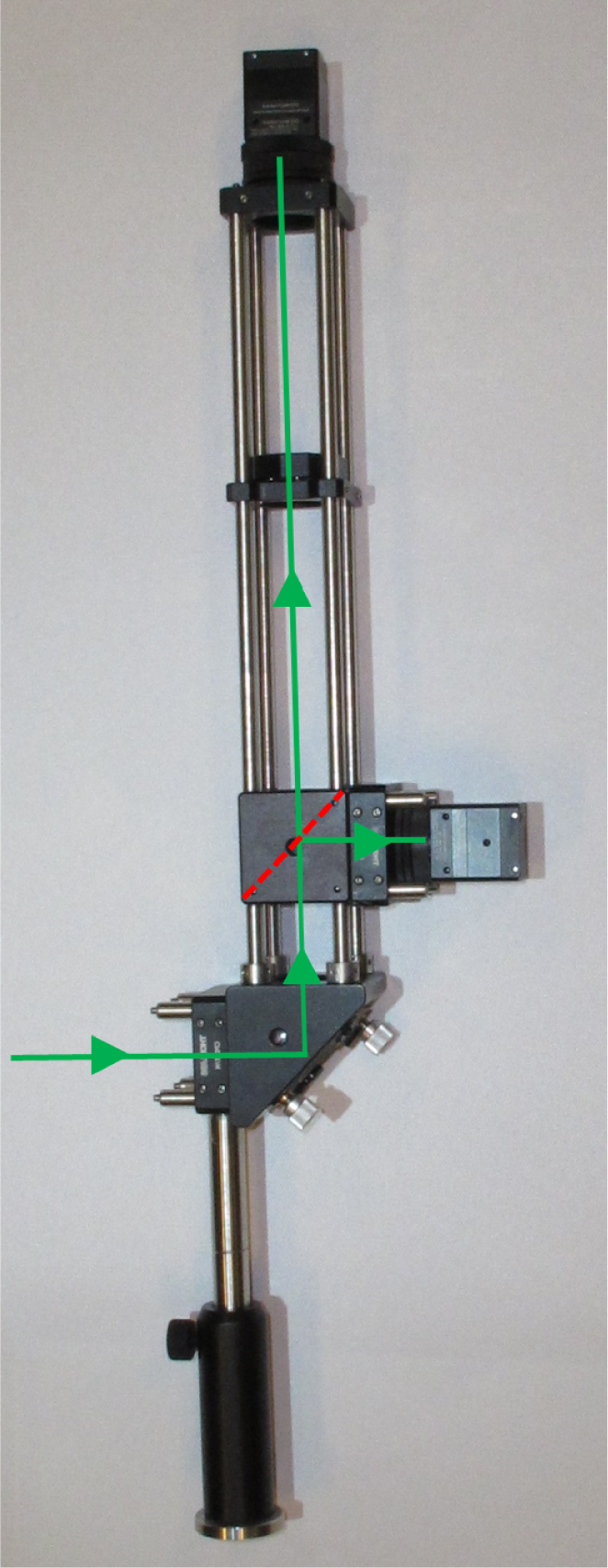
The basic hardware. Diagram showing input beam (green) reflected by 45° mirror and then split at a 50:50 beamsplitter and observed with two cameras.

The addition of a lens in the “long arm” of the dual camera configuration at its focal length from the camera sensor enables an additional configuration. In this additional configuration, if the system is placed in an infinity segment of the beam path, it will simultaneously visualise both the Fourier plane and the image plane. Using a removable kinematic mount, the lens can easily be placed in or out of the beam, quickly converting between this application and the standard dual camera mode.

#### Cameras

Due to their small physical size, high pixel count, and support in the Python Microscope package, we opted to use the Ximea xIQ cameras, although other cameras can easily be used. The Ximea xIQ technical specifications can be found on the vendor’s website (https://www.ximea.com/downloads/usb3/manuals/xiq_technical_manual.pdf).

The pixel size of the Ximea xiQ MQ042MG-CM is 5.5 µm so the lateral optical resolution for both cameras is 11 µm. However, by localising the centroid of the signal above background we get localisation precision significantly better than the image resolution [4] (Figure 2). The exact precision depends on both the spot size and its signal to noise ratio, but simulations suggest that realistic values will generate precisions 10–100 times better than the human eye, ≈0.01 pixel size i.e. 55nm for the Ximea xiQ (Figure 2). The optical angular resolution of the top and bottom cameras are 7.70 and 19.72 arcseconds respectively, but again, the centroid calculation enables localisation of the spot centre to a much higher precision.

**Figure 2:**
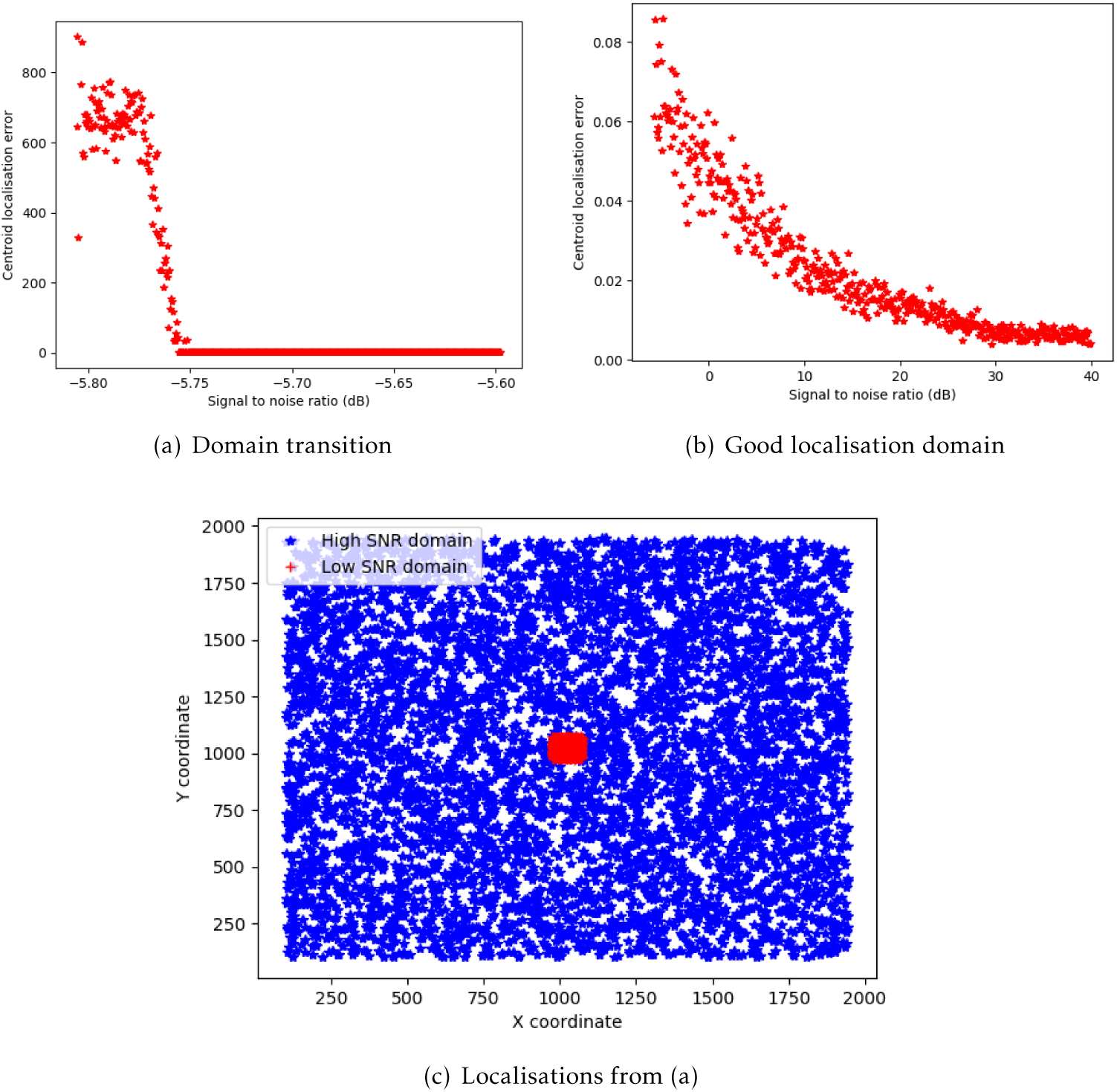
Measuring the localisation accuracy as a function of signal to noise ratio (SNR) using simulated data. A Gaussian spot 200×200 pixels with standard deviation of 40 pixels was placed in a known random location in a 2048×2048 array with Gaussian white noise at a fixed SNR. The centre was measured using the same process as BeamDelta. (a) Centroid localisation error as a function of SNR, centred around the domain transition threshold between poor and good localisation. The “transition zone” spans −5.78 dB to −5.76 dB. (b) Centroid localisation error as a function of SNR in the good localisation domain. For normal use case SNR, (20–40 dB i.e. SNR 10:1-100:1) the localisation accuracy is ≈0.01 pixels. (c) A visual representation of the localisations in (a). In the low SNR domain, the centroid is always placed at roughly the centre of the array. In the high SNR domain, the centroid position varies randomly as expected since the position of the Gaussian spot varies randomly.

The higher angular precision of the top camera implies that it should be used for measuring any angular deflection. Although the human eye has an theoretical angular resolution of 3–6 arcseconds, for reasons discussed in the Introduction, it is unlikely that this is ever achieved in practice. The centroid measurements presented here can achieve significantly better precision even at low signal to noise ratios (Figure 2). It is therefore clear that BeamDelta offers a distinct advantage over conventional alignment practises using visual observation of beams on pinholes or irises.

#### Software

The software component of BeamDelta provides a live view of the cameras, the reference and current centre positions of the brightest feature on the camera, usually a laser beam spot, and the distance between these two positions (Figure 3).

**Figure 3:**
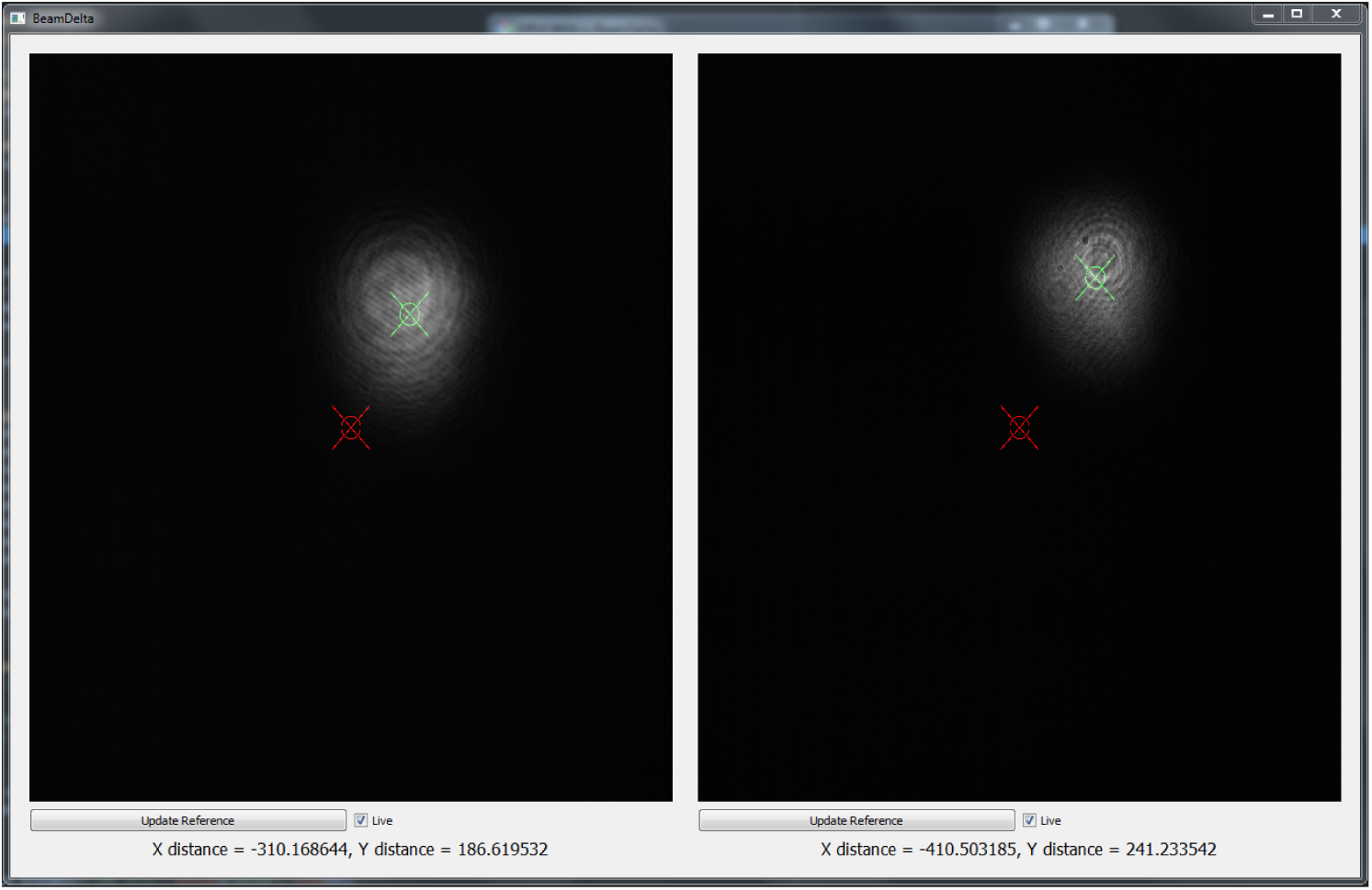
Screenshot of the BeamDelta user interface. The red cross marks the location of the centroid to be aligned to (the “reference centroid”). The green cross marks the location of the current centroid. The difference in X-Y positions of the centroids for each camera is shown underneath as “X distance” and “Y distance”. The “Live” tick box sets whether the BeamDelta is collecting current data from each camera. Clicking “Update Reference” sets the location of the reference centroid to be the same as the location of the current centroid.

The software is implemented in the Python programming language and uses the Qt widget toolkit to create a simple graphical user interface (GUI), the Python Microscope package to control the cameras, and SciPy and scikit-image to measure the laser beam centre.

For each camera, BeamDelta has an `AlignmentControl` instance. This class represents the current alignment of a laser beam on its corresponding camera. On the GUI, each `AlignmentControl` has a corresponding visual and text view. The visual view shows the current camera image with two marks, the current laser beam centre and the reference position. The text view shows the distance between the current and reference positions in pixels.

The centre positions are measured by computing the centre of mass, or weighted centroid, of the laser beam image region. The laser beam image region is segmented from background using Otsu’s threshold algorithm [6].

When a camera acquires a new image, the `AlignmentControl` instance computes the centre position of the laser beam and its offset from the reference position. As alignment changes, the visual and text views update their display.

### Operation

#### Setup

The BeamDelta software is available on the Python Package Index (PyPI). `pip` is the Python recommended package installer and handles BeamDelta dependencies automatically. Therefore, the easiest method to install BeamDelta is:

~~~
pip install BeamDelta
~~~

BeamDelta connects to Python Microscope device servers, one for each camera. The instructions on how to do this are specific to the cameras being used and are part of the Python Microscope documentation. In brief, the Python Microscope `deviceserver` program will create a device server for each camera and display a corresponding URI which has a format like:

~~~
PYRO:[microscope_device_name]@[ip_address]:[port]
~~~

Once one, two, or more camera servers have started and their URIs are known, the BeamDelta program can be started with:

~~~
BeamDelta CAMERA1-URI [CAMERA2-URI […]]
~~~

BeamDelta starts capturing live images from the cameras. The reference position, i.e. the position which the laser beam is being aligned to, is displayed as a red cross and starts at the centre of the camera display. The current beam centre is displayed as a green cross. A snapshot of this UI is show in Figure 3.

At this point, the BeamDelta hardware should be added to the path and roughly aligned so that the beam images fall on the sensors. The exact BeamDelta hardware configuration is dependent on the usage purpose and is detailed on the Hardware section.

While the BeamDelta hardware does not need to be well aligned, it is helpful to have the reference position in the centre of the camera views for two reasons. First, since the beam being aligned starts with an offset from the reference position, if the reference position is near the edges of the camera then the beam being aligned is more likely to start outside the camera view. Second, computation of the beam centre is dependent on measuring the whole beam spot, so if the beam is closer to edge, parts of it may be clipped. We generally consider a reference centroid calculated within the central 30% of the camera’s display to be sufficent for most practical alignment purposes but this is application dependent, as large spots compared to the camera sensor size will start to be clipped at the edge with small deflections from the centre of the sensor.

Once the BeamDelta hardware is in place, the reference beam position needs to be updated by clicking on the “Update Reference” button.

#### Use Case 1: Lens Alignment

In the first use case, a lens is being introduced into a beam path which has already been aligned to a satisfactory degree. Typically the lens is mounted in a lens holder with XY-position manipulators. BeamDelta is placed in the beam path before the lens is added and the reference position without the lens is recorded by pressing “Update Reference” button. The lens is then introduced and approximately positioned so that the beam passes near the centre of the lens and the lens is roughly perpendicular to the beam path. BeamDelta will measure the current deflection introduced to the beam path by the lens. The XY position of the lens should then be adjusted until the position of the current centroid matches the position of the reference centroid. The back reflection is then checked to see if it overlaps with the input beam. This is adjusted by rotating the lens. The alignment steps are then repeated until the offset is sufficiently small (this is obviously application dependant, but we usually us a threshold of <1 pixel, in BeamDelta) and the back reflection overlaps the incident beam. Figure 4 shows BeamDelta in use aligning a lens in a bespoke microscope system. This use case can be achieved using either the Single Camera or Dual Camera configurations since the lens only alters the position of the beam.

**Figure 4:**
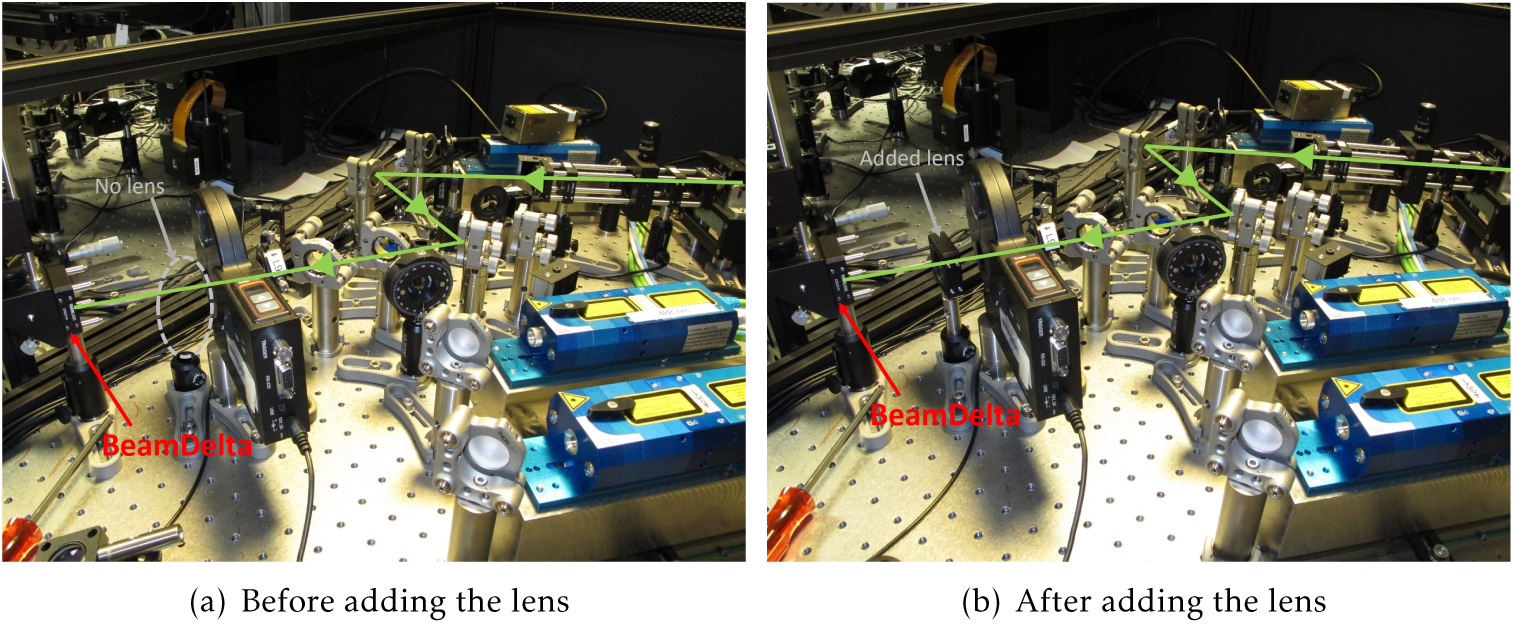
Lens Alignment use case: An aligned beam path terminating at BeamDelta is measured and the position of the beam centroid marked as reference. A lens is then added to the beam path and aligned to the reference to ensure that the beam position after the lens is added is the same as the position before the lens was added. (a) Aligned beam path before the lens is added (b) Beam path after the lens is added.

#### Use Case 2: Laser Co-alignment

In the second use case, one or more lasers are being aligned to an existing beam path. In this case, the additional beam must have some mechansim by which its angle and position can be adjusted. Typically, this is achieved by having the new beam reflect off two mirrors before being directed into joining the existing beam path. One of these mirrors can be the dichroic used to merge the beams, and this is the setup we use. BeamDelta is placed at any suitable point in the existing beam path and the reference position calculated from the current position of the beam on both cameras, generating a reference path. Then, the existing laser is turned off and the new laser is turned on. BeamDelta will measure the misalignment between the new beam path and the existing beam path as the difference between the current and reference centroid positions. The mirror closer to the laser is used to adjust the position of the current centroid on the lower camera display such that is overlaps with the reference position. The mirror further from the laser is used to adjust the position of the current centroid on the upper camera display in a similar way. Doing this will cause the position of the lower camera centroid to be deflected again, although it should be by a lesser degree. By iteratively correcting for each camera using the corresponding mirror, eventually the position of both will be co-aligned with the reference positions. At this point, the new beam path will be co-aligned with the existing beam path. Figure 5 shows BeamDelta in use co-aligning a 488 nm laser to a pre-aligned 561 nm laser in a bespoke microscope system. Since this use case requires alignment of both the position and angle of the new laser, the Dual Camera configuration is necessary.

**Figure 5:**
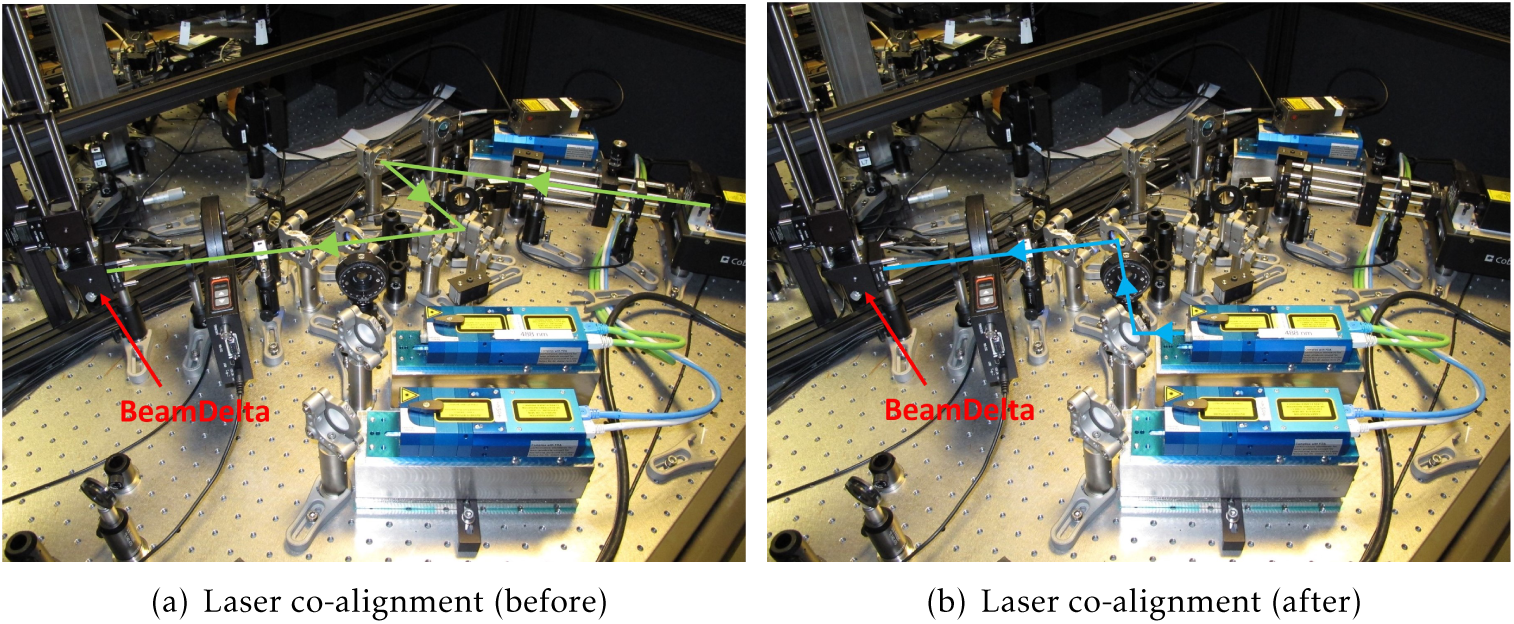
Laser Co-alignment use case: An aligned beam path terminating at BeamDelta is measured and the position of the beam centroid marked as reference. The first laser is turned off and a second laser turned on. The difference in position is measured and the second laser is aligned to the position and angle of the first. (a) Initial aligned beam path. (b) Second beam path after co-alignment.

#### Requirements

The hardware requirements for BeamDelta are rather modest and flexible. It isn’t necessary for the two cameras being used in the setup to have the same pixel size or sensor size. We recommend that, in the relevant configurations, the upper and lower cameras should have sufficiently different path lengths as this determines the sensitivity to angular divergence between beams. We typically use a path difference of more than 150 mm. Shorter path differences will reduce angular sensitivity. The main hardware requirement for BeamDelta is that it fits in the optical setup. If the Dual Camera configuration is mounted parallel to the optical axis, the space required for both arms may prove problematic. If the hardware is mounted perpendicular to the optical axis using a 45° plane mirror as we show in Figure 1, then there only has to be space for the aforementioned plane mirror making it far easier to drop BeamDelta at various point in the beam path for alignment.

The BeamDelta program uses the Python Microscope package to interface to the cameras and so is limited to the cameras supported by Microscope. Microscope is under active development so refer to its documentation for the list of supported cameras. Currently, the supported cameras are Ximea, Andor (SDK2 and SDK3), and Photometrics/Roper/Qimaging cameras using pvcam.

In terms of software, the BeamDelta program is computationally undemanding and will run in virtually any computer. It is written in the Python programming language and requires the Python packages Microscope, NumPy, PyQt5, SciPy, and scikit-image, all of which are free and open source software. We have tested BeamDelta in GNU/Linux, MacOS, and Windows operating systems.

In addition to the BeamDelta program requirements, the choice of camera may add specific requirements. For example, Ximea cameras require the XIMEA Application Programming Interface which is non-free software and is available for selected GNU/Linux, MacOS, and Windows versions. Another example are cameras that communicate over Camera Link, requiring a PCI/PCIe card which means the computer will need a PCI/PCIe slot. Refer to Python Microscope (https://www.python-microscope.org/) and the camera vendor documentation for the specific requirements of each camera.

## Discussion

BeamDelta is a tool to assist with alignment of optical systems. It shows the difference between the current beam path and a reference path. As such, it can be used to align optical components by returning a beam path to its original position after introduction of an optical component, or to align one beam against another. It is important to note that BeamDelta provides relative alignment information, and its use depends upon starting with a reference position for relative alignment to.

As mentioned previously, a significant strength of BeamDelta as an alignment tool is that it doesn’t require a particularly rigorous alignment in order to be useful. However, it is worth noting that the position of the reference point is important. If this is significantly off centre in any particular direction in one or both of the cameras, the amount of lateral or angular deflection that can be measured along that direction before some part of the beam falls off the camera sensor will be less.

As previously mentioned, the localisation precision of BeamDelta offers a significant improvement over previous methods of alignment, with the human eye having an accuracy of ≈10 µm given reasonable assumptions. We estimate that a realistic use case will enable ≈55 nm lateral resolution and similar improvements in angular resolution. It also offers accurate recall of both previous beam locations and the difference between current and previous beam locations. The localisation accuracy is at least an order of magnitude better than that possible by eye alone, once a minimum threshold signal to noise ratio is exceeded. In the extremely low signal to noise ratio (SNR) domain, the Otsu thresholding fails and the localisation consistently defaults to the centre. At signal to noise ratios above ≈−5.77 dB the localisation accuracy rapidly becomes sub-pixel. The SNR of −5.77 dB represents a signal to noise root mean square (RMS) ratio of 1:0.51 since *SNR*_dB_ = 10 log_10_((*Signal*_*RMS*_ */Noise*_*RMS*_)^2^). All the use cases envisioned for BeamDelta involve direct launching lasers into the cameras and the main noise sources will be ambient room lighting and read noise. It is therefore extremely unlikely that a user is ever going to observe the low SNR domain and so no other thresholding options, such as manual thresholding, have been implemented.

The BeamDelta software is available as a Python package under a free and open source license. We suggest that it is installed with the Python package manager pip as this will install the required Python dependencies. Additionally, specific camera hardware drivers may be required. Once installed the cameras must be configured within Microscope and their addresses noted to add to the BeamDelta command line.

Future improvements include automated detection of cameras, and audible feedback with a rising tone indicating improving alignment.

BeamDelta is a powerful tool for aiding those working with bespoke optical paths, such as microscopes. It offers significantly higher accuracy and repeatability over the established technique of visual observation of beams on pinholes or irises. Additionally, it eliminates the effect of parallax errors in alignment since the alignment measurements are made in line with the optical path. Using BeamDelta also offers significant improvements in safety as, accurate alignment can be performed without the use of handheld viewing aids or the removal of protective eyewear. Overall, BeamDelta offers significant improvements over traditional alignment methods.

## Software availability

1. BeamDelta code repository: https://github.com/MicronOxford/BeamDelta
2. Software license: GNU General Public License version 3 or any later version (https://www.gnu.org/licenses/gpl-3.0.en.html)

## Acknowledgements

We would like to thank Micron Oxford, Martin Booth, Mick Philips, Mantas Žurauskas, and Jingyu Wang for helpful comments and suggestions during the development of BeamDelta.

